# Leaf δ^13^C reveals post-photosynthetic fractionation during ontogeny in a C_4_ grass

**DOI:** 10.1101/2023.09.25.559424

**Authors:** Yong Zhi Yu, Hai Tao Liu, Fang Yang, Lei Li, Rudi Schäufele, Guillaume Tcherkez, Hans Schnyder, Xiao Ying Gong

## Abstract

The ^13^C isotope composition (δ^13^C) of leaf dry matter is a useful tool for physiological and ecological studies. However, how post-photosynthetic fractionation associated with respiration and carbon export influences δ^13^C remains uncertain. We investigated the effects of post-photosynthetic fractionation on δ^13^C of mature leaves of *Cleistogenes squarrosa*, a perennial C_4_ grass, in controlled experiments with different levels of vapour pressure deficit and nitrogen supply. With the increase of leaf age classes, the ^12^C/^13^C fractionation of leaf organic matter relative to the δ^13^C of atmosphere CO_2_ (Δ_DM_) increased while that of cellulose (Δ_cel_) was almost constant. The divergence between Δ_DM_ and Δ_cel_ increased with leaf age classes with a maximum value of 1.6‰, indicating the accumulation post-photosynthetic fractionation. Applying a new mass balance model that accounts for respiration and export of photosynthates, we found an apparent ^12^C/^13^C fractionation associated with carbon export of –0.5 to –1.0‰. Different Δ_DM_ among leaves, pseudostems, daughter tillers and roots indicate that post-photosynthetic fractionation happens at the whole-plant level. Compared with Δ_DM_ of old leaves, Δ_DM_ of young leaves and Δ_cel_ are more reliable proxies for predicting physiological parameters due to the smaller sensitivity to post-photosynthetic fractionation and the similar sensitivity in responses to environmental changes.

**BRIEF SUMMARY STATEMENT:** Δ^13^C of bulk organic matter increases with leaf age classes while Δ^13^C of cellulose remain constant, lending support to the use of Δ^13^C of cellulose as a more reliable proxy for predicting physiological parameters due to the smaller sensitivity to post-photosynthetic fractionation.

## INTRODUCTION

The carbon isotope composition (δ^13^C) of plant organic matter is an important proxy for physiological status of plants. Photosynthetic ^12^C/^13^C discrimination (Δ) is directly related to the ratio of CO_2_ mole fractions in the intercellular space to that of the atmosphere (*C*_i_/*C*_a_), thus can be used to estimate intrinsic water use efficiency (iWUE, Farquhar, O’Leary & Berry 1982, von Caemmerer *et al*. 2014; Gong *et al*. 2023). Δ of C_4_ species is associated with the efficiency of the CO_2_ concentrating mechanism, influenced by *C*_i_/*C*_a_ and bundle sheath leakiness (*Φ*, the ratio of CO_2_ leak from bundle-sheath cells to the rate of PEPc carboxylation) (Farquhar 1983; von Caemmerer *et al*. 2014; Gong, Schäufele & Schnyder 2017), providing valuable information of photosynthetic properties of C_4_ species. Understanding Δ of C_4_ species is essential for inferring the terrestrial carbon isotope signature (Still *et al*. 2003) or for estimating contributions of C_3_ vs. C_4_ plants to terrestrial vegetation or soil organic carbon (Wittmer *et al*. 2010).

Δ is typically assessed by biomass-based discrimination (Δ_DM_), which is a time-integrated proxy for photosynthetic discrimination (Wang *et al*. 2012; Meinzer & Zhu 1998; Meinzer, Plaut & Saliendra 1994; Fravolini Williams & Thompson 2002). However, the discrepancy between photosynthetic and biomass-based discrimination leads to errors in estimating physiological parameters (e.g., *Φ* and *C*_i_/*C*_a_) when Δ_DM_ is directly interpreted as photosynthetic discrimination (Kubásek *et al*. 2007; Kromdijk *et al*. 2014). Across C_4_ species (including grasses and dicots), the discrepancy between Δ assessed by dry matter and the combined gas exchange and Δ measurements (Δ_A_) ranged 0.7-2.4‰, corresponding to a difference in estimated *Φ* of *c*. 0.15 (Kubásek *et al*. 2007). This discrepancy is not trivial because Δ of C_4_ species is typically below 8‰ (Cornwell *et al*. 2017) and considered less sensitive to environmental stimuli compared to that of C_3_ species (von Caemmerer *et al*. 2014; Cernusak *et al*. 2013; Henderson, Caemmerer & Farquhar 1992). In controlled environmental studies, the variation of Δ_A_ of C_4_ leaves was typically smaller than 2‰ in response to nitrogen nutrition or vapour pressure deficit (VPD) (Gong, Schäufele & Schnyder 2017; Meinzer & Zhu 1998).

The difference between photosynthetic and biomass-based Δ is likely related to post-photosynthetic fractionations (Δ_post_) (Badeck *et al*. 2005; Cernusak *et al*. 2009; Cernusak *et al*. 2013; Bowling, Pataki & Randerson 2008). The mixing signals of photosynthetic and post-photosynthetic fractionations are integrated into Δ_DM_, and this mechanism is also contributed to the difference in Δ between heterotrophic and autotrophic (e.g., leaves) organs (Badeck *et al*. 2005; Cernusak *et al*. 2009). Furthermore, many studies have shown a decreasing pattern in leaf δ^13^C during leaf development. δ^13^C of leaves decreased by about 1‰ during early maize ontogeny (Ghashghaie *et al*. 2016). For C_3_ plants, Vogado *et al*. (2020) found a Δ^13^C difference of 1-2 ‰ between expanding and mature leaves of deciduous and evergreen species, which was explained by the changes in leaf photosynthetic δ^13^C and *C*_i_/*C*_a_ during leaf expansion. Obviously, Δ_post_ is not negligible since it can be as large as (or larger than) the environmental effects on Δ_DM_. However, the mechanisms controlling Δ_post_ during ontogenetic development are still poorly understood due to the multitude of metabolic processes (Tcherkez *et al*. 2003; Gilbert *et al*. 2011, 2012).

Respiratory fractionation is recognized as an important mechanism for post-photosynthetic fractionation. Tcherkez *et al*. (2003) demonstrated that the δ^13^C of dark-respired CO_2_ depends on the relative contributions of pyruvate dehydrogenase (PDH) reaction and the TCA cycle. Compared with leaf organic matter, the respiratory CO_2_ is enriched (by *c*. 6.5‰) when PDH activity predominates only, because of the decarboxylation of ^13^C-enriched C atoms of pyruvate, while relatively depleted respired CO_2_ (by –0.5‰) released from the TCA cycle originates from the ^13^C-depleted C atoms in glucose (Rossmann, Butzenlechner & Schmidt 1991; Tcherkez *et al*. 2003). Due to the higher contribution of PDH reaction relative to TCA cycle to the overall respiration, leaf respiration is mostly enriched relative to the respiratory substrates (Brüggemann *et al*. 2011; Ghashghaie *et al*. 2003). Indeed, studies showed that respiratory CO_2_ is enriched by 0∼5‰ relative to the putative substrates in C_4_ leaves (Rooney 1988; Scrimgeour Bennet & Connacher 1988; Lin & Ehleringer 1997; Ghashghaie *et al*. 2016; Sun *et al*. 2012) and 0∼6‰ in C_3_ leaves (Tcherkez *et al*. 2003; Klumpp et al. 2005; Wingate *et al*. 2007; Ghashghaie *et al*. 2001). Another mechanism of post-photosynthetic fractionation is related to the export of assimilates from source leaves. Although transport process itself (including loading, phloem movement, and unloading) does not fractionate (Cernusak, Farqhuar & Pate 2005; Maunoury-Danger *et al*. 2009), biochemical fractionation related to accumulation or loss (e.g., by export) of ^13^C-enriched or -depleted compounds may cause a temporal change of Δ_DM_ of a certain organ (e.g., leaf). Soluble sugars and cellulose are slightly enriched in ^13^C than bulk leaf and lipids (Zhong *et al*. 2017; Ghashghaie *et al*. 2016; von Caemmerer *et al*. 2014). Phloem sap sugars have been reported to be more enriched in ^13^C than the soluble sugars of source leaves (Bögelein, Lehmann & Thomas 2019). Accordingly, the export of ^13^C-enriched sugars out of source leaves can lead to the ^13^C-enrichment in carbon sinks (Bögelein, Lehmann & Thomas 2019; Gessler *et al*. 2008), and ^13^C-depleted lipids and lignin could remain in the leaf (Brüggemann *et al*. 2011; Hobbie & Werner 2004). Based on the conservation of mass, respiration and export-associated metabolism could lead to a ^12^C/^13^C fractionation, which can in turn influence the Δ_DM_ of leaves and result in a trend in Δ_DM_ of successive leaves on plant tillers, although experimental evidence is scarce.

Furthermore, leaf δ^13^C reflects the mixing of source carbon of different origins, that is, C imported from source leaves during the heterotrophic phase and C fixed by the leaf own photosynthesis (Francey *et al*. 1985; Vogado *et al*. 2020). To understand Δ_DM_ of successive leaves, isotopic mass-balance should be considered separately for immature and mature stages. The growth-and-differentiation zone of grass leaves is enclosed within a tube-like structure formed by the sheaths of older leaves (see Fig. 1 in Liu *et al*. 2017a), implying a mostly heterotrophic nature of that tissue. Therefore, grass tillers provide an excellent model system to study post-photosynthetic fractionation processes. For an immature grass leaf, its growth depends on carbon (e.g., water-soluble carbohydrates and proteins) imported from mature leaves (Allard & Nelson 1991) (Fig. 1a). For a mature leaf, carbon import is negligible, and respiration and export are mainly sustained by photosynthates and stored carbon reserves (Fig. 1b).

**Fig. 1.**
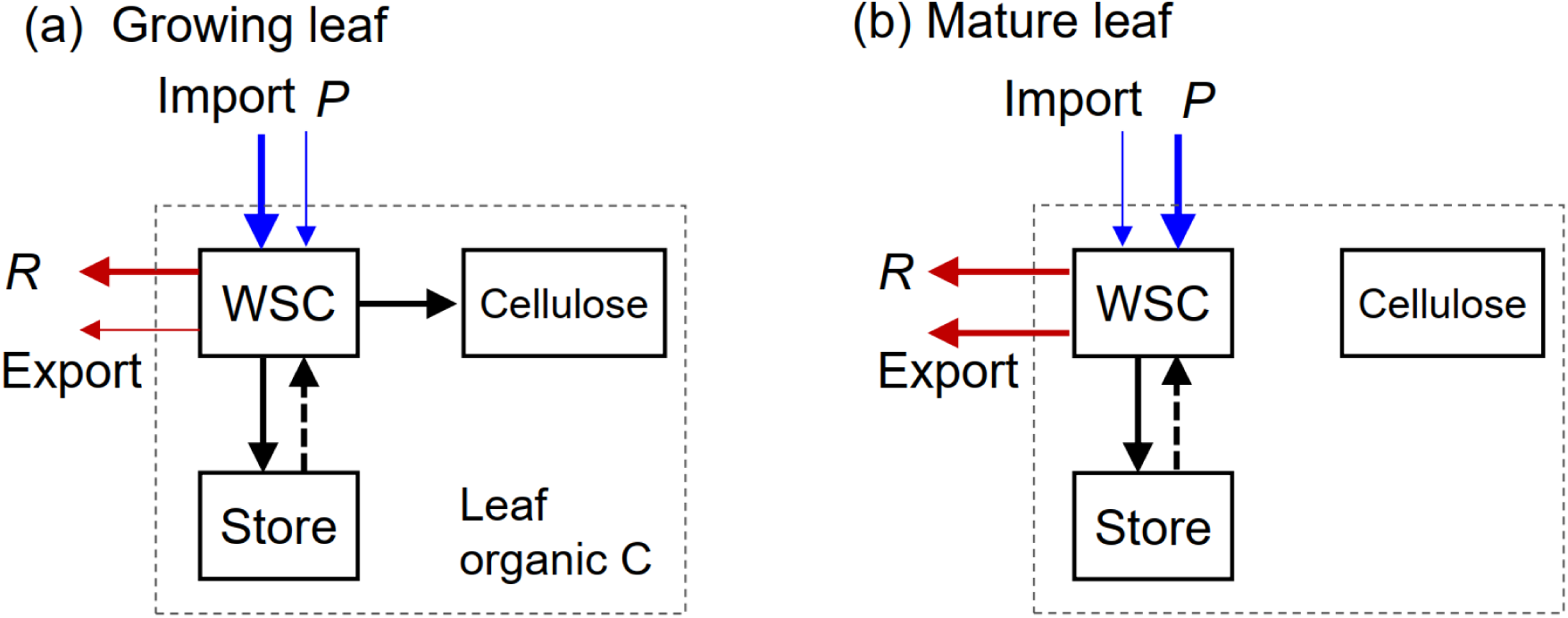
Overview of the processes influencing the isotope composition of leaf organic C of a growing leaf (a) or a mature leaf (b) of grass species. The bulk leaf organic C comprises different pools: water soluble carbon (WSC), cellulose, store, and the rest of C (unspecified in the figure). Arrows indicate C fluxes: import, net photosynthesis (*P*), respiration (*R*), export, and transport between C pools; blue arrows are influxes and red arrows are effluxes.

We took advantage of the sequential nature of leaf development in grasses to evaluate post-photosynthetic fractionations in fully expanded leaves of different ages growing under constant environmental conditions (including stable δ^13^C_CO2_ of the growth atmosphere and irradiance at the top of the canopy). In effect, we achieved a nearly perfectly constant photosynthetic δ^13^C (fixed carbon) of grass canopies to sustain plant growth. This allowed us to rule out the potential influence of unforeseen variation in source carbon δ^13^C. We measured Δ of leaf dry matter (Δ_DM_) and cellulose (Δ_cel_) of mature leaf blades covering an age category of more than 9 ranks on mature tillers of *Cleistogenes squarrosa* (Trin.) Keng. This species is a perennial C_4_ (NAD-ME) grass and is co-dominant in the semiarid steppe of Inner Mongolia. Plants were grown at combinations of moderate or high level of N supply and low or high level of VPD. Half of the plants were grown with ^13^C enriched CO_2_ (δ_CO2_ of –5‰, i.e., 3‰ enriched compared to atmospheric CO_2_) while the other half was grown under a ^13^C depleted atmosphere (industrial CO_2_ with a δ_CO2_ value of –48‰). The consistency in Δ across the two δ_CO2_ conditions should provide strong evidence to confirm results and thus rule out the influence of experimental artefacts (Schnyder *et al*. 2003). We also measured Δ of dry matter of pseudostems of mature tillers, roots, and daughter tillers to evaluate plant-level ^13^C distribution. Theoretically, Δ_cel_ is not affected by post-photosynthetic fractionations after the deposition of cellulose (Liu *et al*. 2017b), thus provides a useful proxy for dictating the variations in photosynthetic δ^13^C during leaf growth. We test the hypotheses that 1) Δ_DM_ changes along leaf-age category while Δ_cel_ remains constant, given that environmental conditions related to photosynthetic discrimination were maintained near constant during plant development; 2) variation in Δ_DM_ along leaf-age category can be explained by respiratory fractionation and the export of carbon from mature leaves. Based on the obtained changes in Δ_DM_ along age category and fractionation in respiration and carbon export, we established a mass-balance model to predict the Δ_DM_ of leaf blades of different age categories.

## MATERIALS AND METHODS

### Treatments and growth conditions

The experiments were performed in four growth chambers (PGR15, Conviron, Canada) which formed part of the controlled-environment mesocosm system as described by Schnyder *et al*. (2003). Individual plants of *C. squarrosa* were grown from seed in plastic pots filled with quartz sand and placed in the growth chambers at a density of 236 plants m^-2^. The study had a 2 × 2 factorial design, with VPD and N supply the two factors, and two levels for each factor. Combinations of VPD and N levels were moderate N ×low VPD (N1 VPD1), moderate N ×high VPD (N1 VPD2), high N × low VPD (N2 VPD1) or high N × high VPD (N2 VPD2). We performed two experimental runs with the first run accommodating high and low VPD treatments at a high N level and the second run accommodating high and low VPD treatments at a moderate N level. Each replicate consisted of one plant stand of a certain VPD ×N combination in a growth chamber. Of the two replicate chambers of each treatment, one received an influx air that was relatively enriched in ^13^C, while the other received ^13^C depleted CO_2_ (Fig. S1). CO_2_ free, dry air was generated by using a screw compressor and adsorption dryer system and mixed with pure CO_2_ before being supplied to the growth chambers (Schnyder *et al*. 2003). CO_2_ and vapor concentrations in the chambers were measured by an infrared gas analyser (LI-6262, LI-COR Biosciences Inc., Lincoln, USA). CO_2_ concentration in the chamber was maintained at 386 ± 3 (SD) μmol mol^-1^ during the light period, and relative humidity in the chamber was maintained constant throughout the diurnal cycle at either 50% (1.58 kPa VPD) or 80% (0.63 kPa VPD). Air temperature was maintained constant at 25 °C throughout the diurnal cycles in all chambers. Modified Hoagland nutrient solution containing either 7.5 mM N (N1) or 22.5 mM N (N2) in the form of equimolar concentrations of calcium nitrate (Ca(NO_3_)_2_) and potassium nitrate (KNO_3_) were supplied three times per day to plant stands that were assigned to the moderate and the high N treatments, respectively. Environmental conditions other than N nutrition and VPD were the same for all chambers: a 16/8h light/dark cycle with a photosynthetic photon flux density of 800 μmol m^-2^ s^-1^ at the canopy height maintained throughout the experiment by adjusting the height of cool white fluorescent tubes, other conditions were detailed by Liu *et al*. (2016).

### Plant sampling and isotopic analysis for bulk dry matter

Plants were sampled four (N1 treatment) or five times (N2 treatment) at 2-or 3-day intervals after canopy closure. On each sampling date, four plants were sampled from each chamber. In total, 32 plants were sampled for both N1VPD1 and N1VPD2; 40 plants were sampled for both N2VPD1 and N2VPD2. From each plant, two major tillers (the main tiller and another mature tiller of similar size) were excised. Leaf Nr. 1 is the youngest leaf which included partially unexposed leaf blade tissue that was enclosed within the ‘next-older leaves’ sheath and represented the ‘immature tissues’ of a major tiller as described by (Yang *et al*. 2020). Mature (that is fully expanded, green) leaves were numbered consecutively from the youngest (Nr. 2) near the tip of the tiller to the oldest near the base of the tiller. Thus, the number/rank of leaves represents the gradient of age and vertical position within the canopy. Mature leaf blades on the two major tillers were cut off at the ligule, and the two blades of the same leaf number were combined as one sample for further analyses. Leaf sheaths, and the pseudostem section of the two major tillers were collected as one tissue fraction called ‘stem’. The remaining shoot material and roots were collected as separate samples, and these fractions of tissue represent ‘daughter tiller’ and ‘root’ fraction, respectively. All biomass samples were dried in an oven at 60 °C for 48 hours, weighed and ground to fine homogeneous powder with a ball mill, and stored before analysis of isotope composition.

The C content and δ^13^C of plant dry matter samples were determined after combustion in an elemental analyzer (NA 1110, Carlo Erba Instrumentation) interfaced to a continuous-flow isotope ratio mass spectrometer (Delta Plus, Finnigan MAT). Laboratory standards (wheat flour) were measured after every 10 samples. The measurement precision (SD) for laboratory standards was 0.16‰ for δ^13^C and 0.37% for C content (C mass per total mass, as %).

### Calculation of isotope fractionation

The carbon isotope discrimination of plant samples was calculated as:

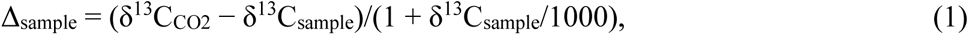

with δ^13^C_CO2_ the CO_2_ in the atmosphere of the growth chamber in which the plant was kept. We calculated carbon isotope discrimination of bulk leaf dry mass (Δ_DM_), daughter tillers (Δ_DT_), stems (Δ_stem_) and roots (Δ_roots_). The δ^13^C of all mature leaf blades on a major tiller (δ_ML_) was calculated as the C mass (C_mass_)-weighted integral of all leaf blade age categories:

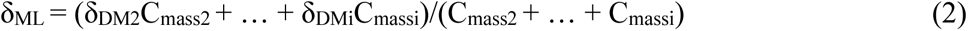

The ^12^C/^13^C discrimination of all mature leaf blades on a major tiller (Δ_ML_) was calculated using Eqn 1 and 2, representing the isotopic signature of source leaves.

### Isotopic analysis for cellulose and mesocosm gas exchange

Additional sets of plants were sampled from all chambers for cellulose extraction and analysis. Three samples of each leaf age category in each chamber were collected by combining leaves of same age category (leaf Nr. 2-10) from 15-20 major tillers (*n*=6 for each age category at each treatment combination). The separation of leaves by age category and the procedure of oven-drying were the same as described above. From these samples, cellulose was extracted using a modified Brendel, Iannetta & Stewart (2000) procedure as detailed in (Liu *et al*. 2016, 2017b). δ^13^C of cellulose samples were measured similarly as that of bulk dry matter samples, and carbon isotope discrimination of cellulose (Δ_cel_) was calculated using Eqn 1.

The controlled-environment mesocosm system measured the concentration and δ^13^C of air stream at the inlet (C_in_, δ_in_) and the outlet (C_out_, δ_out_) of each chamber at a high frequency. During the light period, the δ^13^C of stand-scale net assimilation (δ_A_) was calculated as

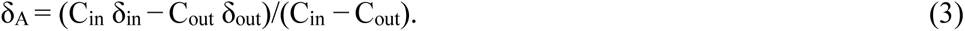

Accordingly, during the dark period, the δ^13^C of stand-scale respiration (δ_R_) was obtained as

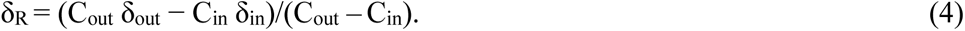

The stand-scale respiratory fractionation was calculated as

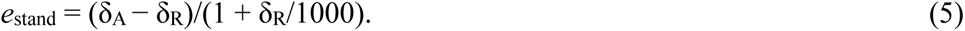

The details of the mesocosm scale gas exchange and isotopic measurements were similar to Schnyder *et al*. (2003) and Gong *et al*. (2017).

### Modelling ^12^C/^13^C discrimination in mature leaves

The leaf organic carbon could be conceptually separated into functionally distinctive pools: water soluble carbon (WSC), transitory storage pool (Store) and cellulose (the main structural carbohydrate). Other major compounds (e.g., protein, lipid, and lignin) were not shown here but included in leaf organic C (Fig. 1). Accordingly, the isotope composition of leaf dry matter (δ_DM_) is an integrated isotopic signature of all carbon pools. δ^13^C of leaf compounds are mainly imprinted from that of import and leaf net photosynthesis (δ_P_). If influx and efflux of carbon carry an isotope composition that differs from δ_P_, δ_DM_ of the leaf will be modified.

For a growing leaf (Fig. 1a) of grasses, carbon is primarily provided by the export of other mature leaves (Allard & Nelson 1991; Turgeon 1989), with a δ^13^C (δ_Imp_) that is similar to the integrated δ_P_ of mature leaves (source leaves). A growing leaf blade is folded and enclosed within the sheathes of older leaves (Liu *et al*. 2017a). Growing leaf also has photosynthetic capacity when the leaf tip is exposed to light (Zhu *et al*. 2020), but its contribution to leaf growth should be marginal. The imported carbon is further used for leaf growth (allocated to structural compounds including cellulose) and metabolism, supporting storage, respiration (*R*) and export.

For a mature (photosynthetically-autotrophic) leaf (Fig. 1b), carbon influx after full elongation is only provided by its own net photosynthesis (*P*, carboxylation rate minus (photo)respiration rate), and respiration in the dark (*R*_D_) and export (*E*_xp_) are the effluxes. According to the mass balance, we have the expression of C mass (C_mass_) of a mature leaf as a function of time-integrated influxes and effluxes:

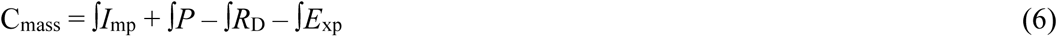

With *I*_mp_ the import of carbon during the growth of the leaf. Applying isotopic mass balance we have,

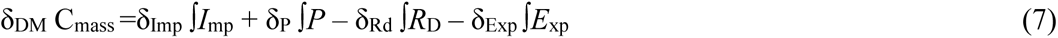

Given that the vast majority of the leaf structural biomass of grass species is inherited from its heterotrophic stage (Robinson-Beers, Sharkey & Evert 1990) and mature leaves of *C. squarrosa* have stopped elongation and their dry mass remained rather constant (Yang *et al*. 2020), C_mass_ of a mature leaf should be close to the time-integrated *I*_mp_, implying that the rate of daily export in mature leaves equal its net assimilation:

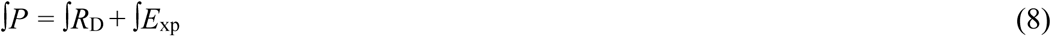

Combing Eqn 7 and 8, we have

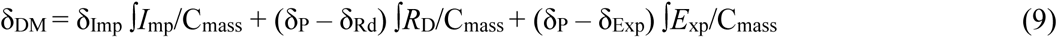

We define the isotopic fractionation of *R* (*e*_resp_) and export (*e*_exp_) relative to photosynthesis (*P*) of a mature leaf as (neglecting second-order terms):

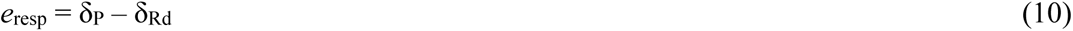

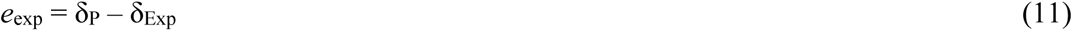

Eqn 9 can be rewritten as:

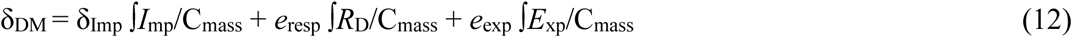

For plants growing in constant environmental conditions, δ_Imp_ was assumed to be constant as the photosynthetic properties of the source leaves remained unchanged. Furthermore, if not, any change of δ_Imp_ would be reflected in the δ of successively synthesized leaf cellulose (δ_cel_), that is cellulose deposited in the successively produced leaves. Cellulose is formed from UDP-glucose, which comes directly from source sucrose with, in principle, little isotope fractionation. In addition, cellulose is mostly deposited during leaf expansion and has no turnover afterwards (Liu *et al*. 2017b). Once the leaf reaches its final length (and is fully expanded) – this developmental step is reflected by the transition from leaf age stages 1 and 2 – δ_cel_ should remain unchanged during aging of the leaf. As will become apparent below, the hypothesis of constant δ_cel_ (and thus δ_Imp_) was verified.

Thus, any change of δ_DM_ of the mature leaf could be explained by isotope fractionation in respiration and metabolism associated with export. The ^12^C/^13^C discrimination of a mature leaf can be calculated as Δ_DM_ = (δ_CO2_ *−* δ_DM_)/(1 + δ_DM_/1000), where δ_CO2_ is the isotopic ratio of the atmosphere. Applying Eqn 12 to the data of leaf Nr. 2 (the youngest mature leaf, i.e., with a both fixed cellulose structural material and functional photosynthesis) and another older leaf (Nr. i) and combining the two equations, Δ_DM_ of leaf i can be calculated from Δ_DM2_ on the same tiller as:

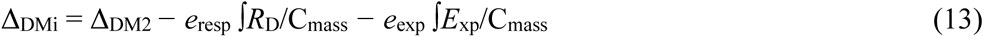

Modelling of Δ_DM_ of individual leaves was performed using Eqn 13, with *e*_resp_ measured from whole-stand gas exchange (Eqn 10), and *R*_D_ was measured by leaf-level gas exchange on the young mature leaves in the same experiments (Gong, Schäufele & Schnyder 2017). The rate of daily export in mature leaves was assumed to equal its net assimilation rate given that all mature leaves has stopped elongation. Export rate was estimated by ^13^CO_2_-labelling of plant for 24 hours (Fig. S2). In brief, steady-state ^13^CO_2_ labelling was performed by swapping randomly selected plants between the two growth chambers of the same treatment combination but supplied with different source CO_2_ (i.e., the ^13^C enriched CO_2_ of –5‰ and the depleted CO_2_ of –48‰), and plants were sampled for isotopic analysis after 24-hour labelling (Schnyder, Ostler & Lehmeier 2017). The daily net assimilation rate in each mature leaf category was estimated as the fraction of new carbon (tracer) in leaf blade:

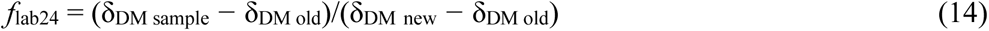

with δ_DM sample_ denotes the isotope composition of the sample, and δ_DM new_ and δ_DM old_ that of control plants, i.e., the end-members of the isotopic mixing model (Gong *et al*. 2014).

The time integrated *R*_D_ or *E*_xp_ was calculated by multiplying *R*_D_ or *E*_xp_ with the age differences between leaf categories which were calculated from the leaf Nr. difference (i.e., i*−*2 in Eqn 13) and a leaf appearance interval of 2.4 days measured in the same experiment (Yang *et al*. 2016). *e*_exp_, the only unknown parameter in Eqn 13, was estimated by using the ‘solver’ function of Excel to minimize the sum of squared error (SSE) of the estimated Δ_DM_ of all leaves for each sampled tiller. Afterwards, the modelled Δ_DM_ of individual mature leaves were plotted against the measured Δ_DM_ and compared with the 1:1 relation. ANOVA analysis was performed to test the effects of N supply, VPD, leaf age category, and their interactions on Δ_DM_ and Δ_cel_ and the effects of N, VPD and their interactions on *e*_resp_, *e*_exp_, export rate and differences in δ^13^C of different tissue fractions using the general linear model of SPSS Statistics 19 (IBM, USA) or “aov” function of R software (version 4.2.1). It should be noted that in Eqn 13, we assume that isotope fractionations *e*_resp_ and *e*_exp_ (Eqns10 and 11), are assumed constant, that is, metabolic properties at the origin of respiratory and export fractionations were assumed to be similar regardless of leaf age in mature leaves. This assumption is probably not critical considering the relatively small *e* values (1 per mil or less). The significance of *e*_resp_ and *e*_exp_ is further discussed below in the section Discussion.

## RESULTS

### Effects of nitrogen, VPD and leaf age category on Δ_DM_ and Δ_cel_

Δ_DM_ increased with increasing leaf age in all treatments (Fig. 2a, b). Δ_DM_ was affected by VPD (*P***<**0.001) and N treatment (*P***<**0.001): Δ_DM_ at high VPD was *c*. 1‰ higher relative to low VPD; plants grown with high N supply had a slightly higher Δ_DM_ than plants with low N supply at both VPD levels (Fig. 2a, b, Table 1). The interaction effect of N supply and leaf number on Δ_DM_ was significant (*P***<**0.001): the slope of the linear regression between Δ_DM_ and leaf number was lower for the low N supply than for the high N supply at both VPD treatments (Fig. 2a, b, Table 1).

**Fig. 2.**
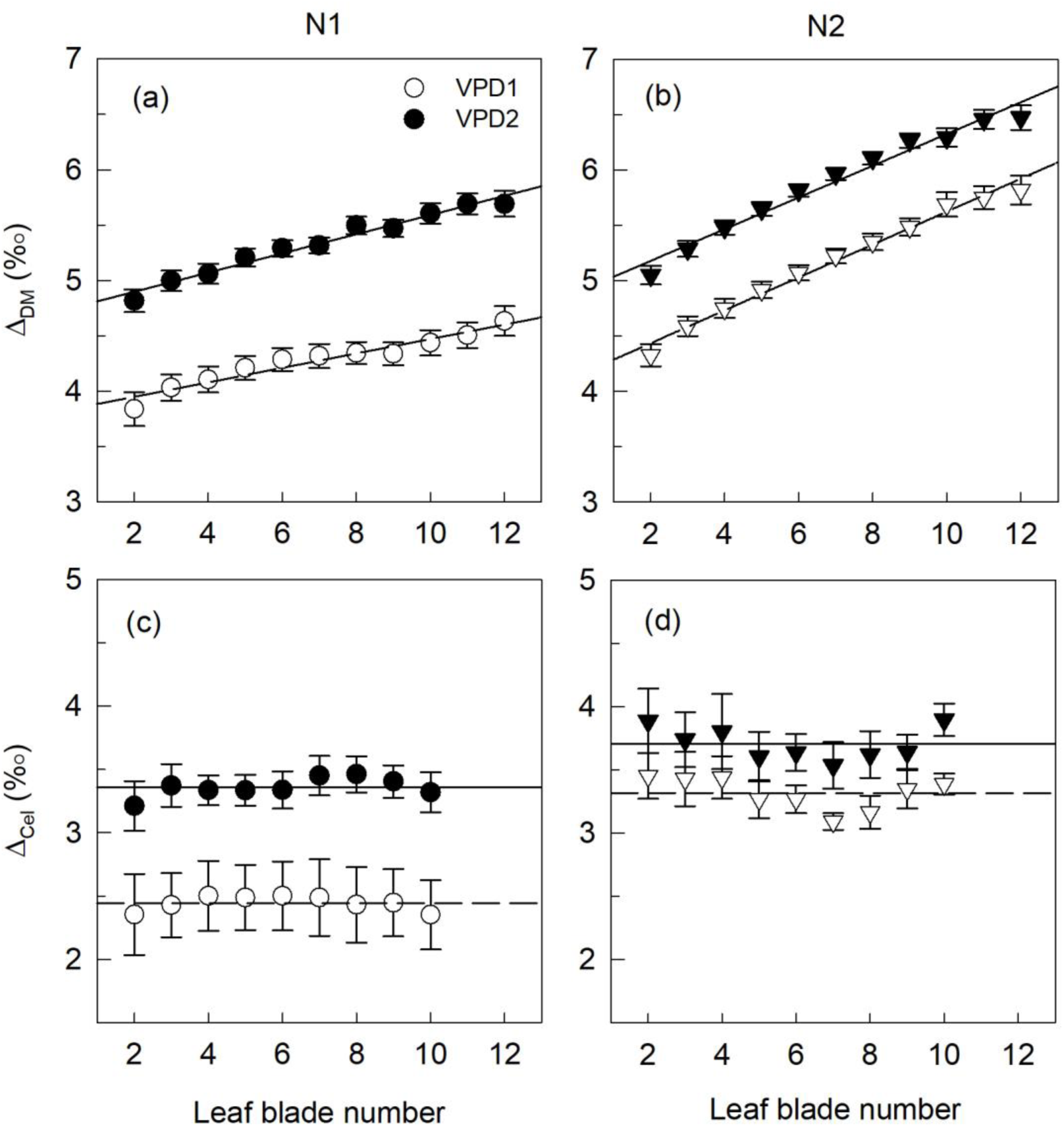
Δ_DM_ and Δ_cel_ of mature blades of a major tiller at moderate or high N supply combined with low or high VPD. Leaves were numbered basipetally, that is from the youngest fully expanded leaf (leaf Nr. 2) to the oldest. Lines indicate linear regression of Δ_DM_ vs. leaf blade number in panel a and b, and means of Δ_cel_ across age categories. Error bars are standard errors (SE), and *n*=18-40 for Δ_DM_ and *n*=6 for Δ_cel_.

**Table 1.** ANOVA table for effects of nitrogen and VPD treatment and leaf age category on Δ_DM_ and Δ_cel_.

| Source | $\Delta_{DM}$ | | | $\Delta_{cel}$ | | |
| --- | --- | --- | --- | --- | --- | --- |
|  | df | *F* | *p* | df | *F* | *p* |
| N | 1 | 697.9 | **<0.001** | 1 | 16.5 | **<0.001** |
| VPD | 1 | 1018.0 | **<0.001** | 1 | 39.9 | **<0.001** |
| N*VPD | 1 | 36.5 | **<0.001** | 1 | 7.4 | **<0.01** |
| Leaf Nr. | 10 | 63.4 | **<0.001** | 8 | 0.8 | 0.59 |
| N*Leaf Nr. | 10 | 6.6 | **<0.001** | 8 | 0.5 | 0.86 |
| VPD*Leaf Nr. | 10 | 0.2 | 0.99 | 8 | 0.6 | 0.74 |

Leaf number had no effects on carbon isotope discrimination calculated from cellulose (Δ_cel_): Δ_cel_ was virtually constant across all leaf ranks (*P*>0.05, Fig. 2c, d, Table 1). Δ_cel_ was higher at high VPD than low VPD and higher at high than low N level (*P***<**0.001, Fig. 2c, d, Table 1). The difference (δ_cel_ − δ_DM_) increased with the increasing leaf number and ranged from 0.8 to 2.6‰ (Fig. 3). N supply and VPD had similar effects on Δ_cel_ and Δ_DM2_: high VPD or high N led to increased Δ_cel_ and Δ_DM2_ (Fig. 4). We observed a mean difference of 1.5‰ between Δ_cel_ and Δ_DM2_, that is, dry matter of young leaves was enriched in ^13^C by 1.5‰ relative to cellulose of the same leaves (Fig. 4).

**Fig. 3.**
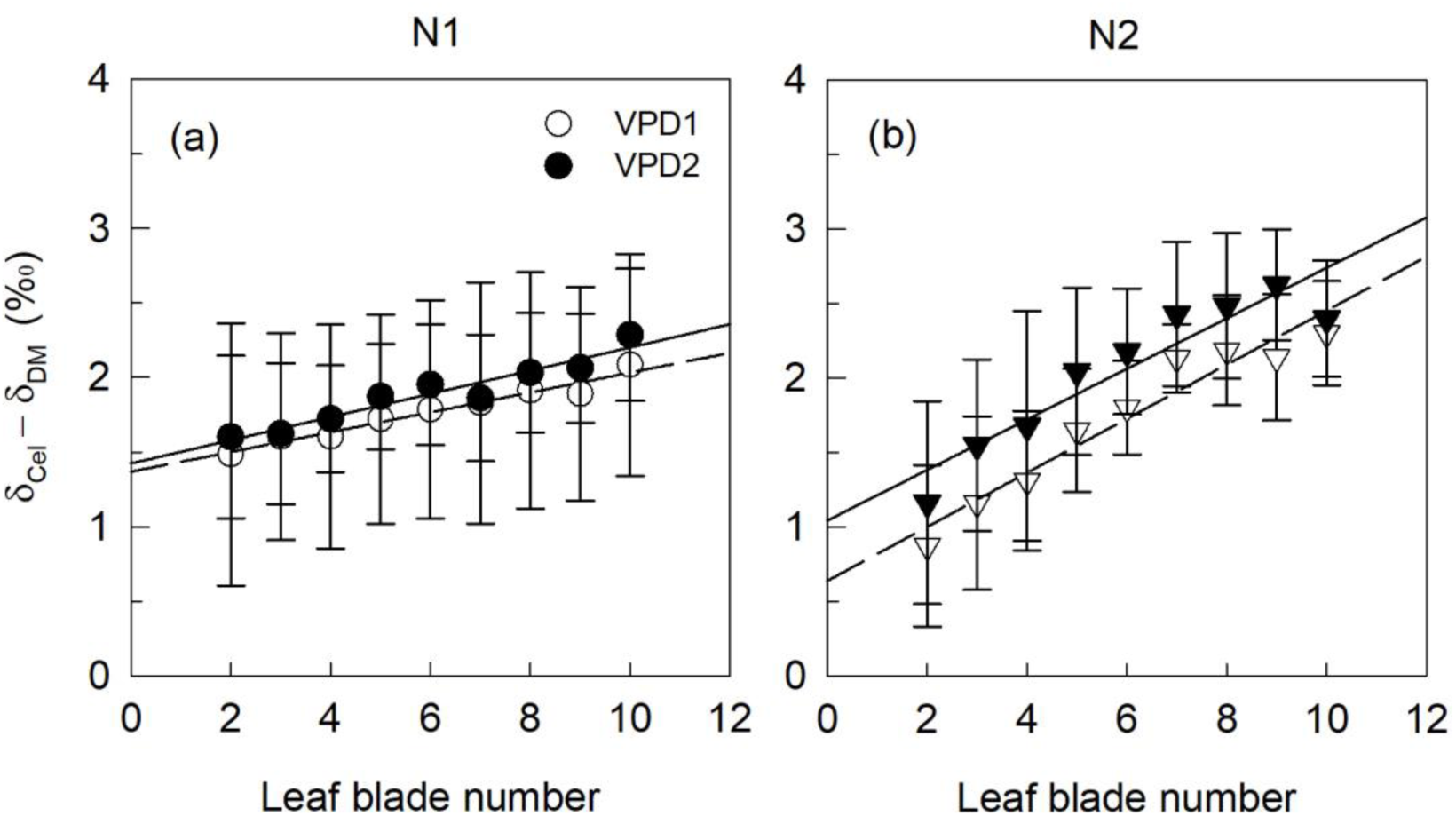
Differences in δ^13^C between cellulose and bulk dry matter (δ_cel_ − δ_DM_) of mature blades of a major tiller at moderate or high N supply combined with low or high VPD. Error bars represent 95% confidence intervals which were calculated by propagating errors in δ_cel_ and δ_DM_.

**Fig. 4.**
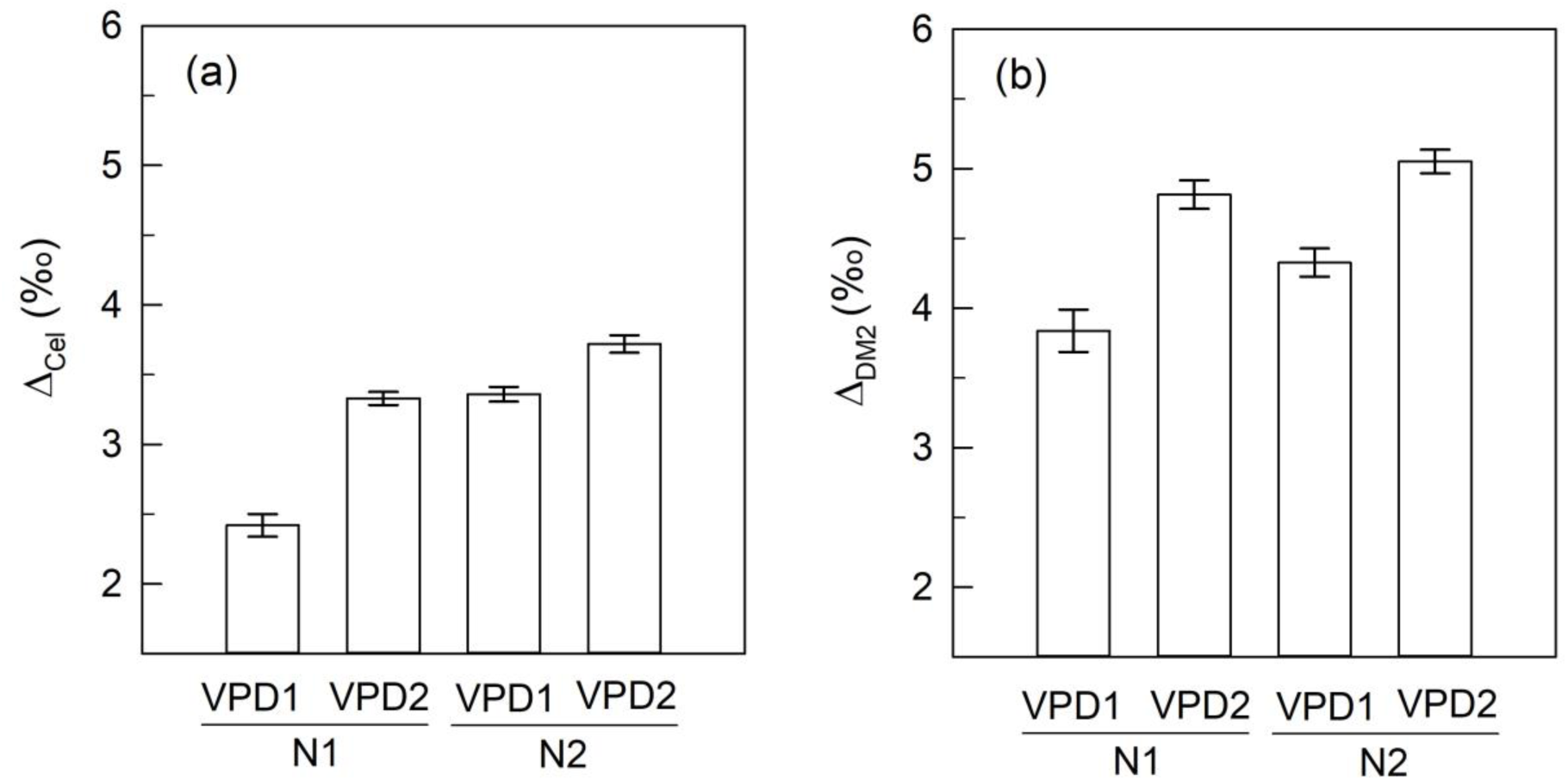
Treatment effects on Δ_cel_ averaged over leaf age categories and Δ_DM2_ (Δ_DM_ of leaf Nr. 2, the youngest fully expanded leaf). Error bars are standard errors (SE), and *n*=6 for Δ_cel_ and *n*=40 for Δ_DM2_.

### Modelling of Δ_DM_

Using Eqn 13, we estimated the isotope fractionation based on dry matter, Δ_DM_, of all mature leaves (Fig. 5). To parameterize the new model, we measured *e*_resp_ and *R*_D_ and obtained export rate from ^13^C-labeling. The only unknown parameter was the apparent isotope fractionation associated with export, *e*_exp_, and was estimated by nonlinear fitting. *e*_resp_ was not influenced by N supply and VPD, with a mean value of –0.9‰ (*P*>0.05, Table 2). The average value of *e*_exp_ across all treatments was relatively small, of –0.68‰, indicating that export metabolism was associated with only a small fractionation in favour of ^13^C. N supply, VPD and their interaction had significant effects on *e*_exp_ (*P***<**0.05, Table 2). N supply increased *e*_exp_ and this effect was stronger at low VPD than at high VPD. The potential influence of parameterisation (export rate and *e*_resp_) on fitted *e*_exp_ is shown in Table S1. There was a relatively low sensitivity of *e*_exp_ to parameterisation, *e*_exp_ values being always in the same order of magnitude. Also, when compared at the same N supply, export rate was higher at high VPD than at low VPD (*P*<0.05, Table 2).

**Fig. 5.**
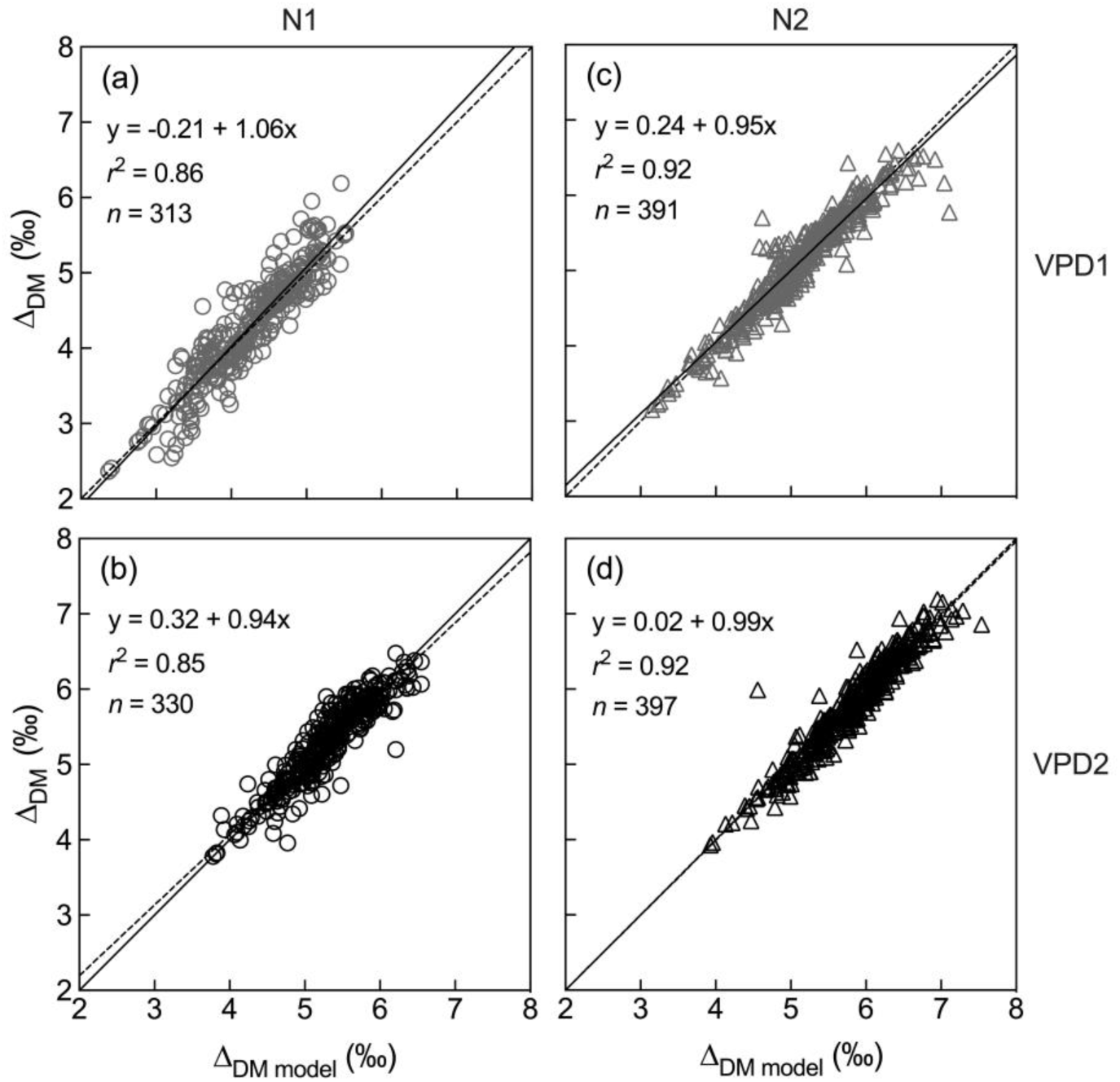
The stimulated Δ_DM_ of individual mature blades using Eqn 13 compared with the measured Δ_DM_ at the combinations of moderate (N1, circles) and high N supply (N2, triangles) and low (grey symbols) and high VPD (black symbols). The dashed line is the 1:1 line for comparison. The black lines are linear regression lines, *p*<0.0001 for all regressions.

**Table 2.** Mean ^12^C/^13^C fractionation (‰): *e*_resp_, stand-scale respiratory fractionation relative to photosynthesis (*n*=4); *e*_exp_, fractionation of carbon export relative to photosynthesis (*n*=8); differences in δ^13^C of different tissue fractions: mature leaves and stems (δ_ML−_ δ_stem_, *n*=32-40), mature leaves and daughter tillers (δ_ML−_ δ_DT_, *n*=32-40), mature leaves and roots (δ_ML−_ δ_root_, *n*=8-21). Export rate (g g^-1^C d^-1^) is equal to *f*_lab24_ calculated from mature leaves with number 2-10 (*n*=4). Mean ±SE are shown.

|  | N1 |  | N2 |  | *p*-value |  |  |
| --- | --- | --- | --- | --- | --- | --- | --- |
|  | VPD1 | VPD2 | VPD1 | VPD2 | N | VPD | N*VPD |
| $e_{\text{resp}}$ | $-1.09 \pm 0.83$ | $-0.75 \pm 0.34$ | $-0.74 \pm 0.51$ | $-1.03 \pm 0.21$ | 0.952 | 0.952 | 0.562 |
| $e_{\text{exp}}$ | $-0.57 \pm 0.06$ | $-0.50 \pm 0.06$ | $-0.96 \pm 0.04$ | $-0.68 \pm 0.04$ | **<0.001** | **<0.01** | **<0.05** |
| $\delta_{\text{ML}} - \delta_{\text{stem}}$ | $-0.03 \pm 0.04$ | $-0.09 \pm 0.05$ | $-0.25 \pm 0.04$ | $-0.32 \pm 0.02$ | **<0.001** | 0.082 | 0.902 |
| $\delta_{\text{ML}} - \delta_{\text{DT}}$ | $0.00 \pm 0.05$ | $-0.14 \pm 0.05$ | $-0.13 \pm 0.04$ | $-0.32 \pm 0.04$ | **<0.001** | **<0.001** | 0.479 |
| $\delta_{\text{ML}} - \delta_{\text{root}}$ | $-0.93 \pm 0.10$ | $-1.09 \pm 0.17$ | $-0.69 \pm 0.12$ | $-0.96 \pm 0.11$ | 0.235 | 0.072 | 0.697 |
| export rate | $0.058 \pm 0.005$ | $0.075 \pm 0.004$ | $0.072 \pm 0.006$ | $0.099 \pm 0.007$ | 0.062 | **<0.05** | 0.588 |

There was a strong positive correlation between stimulated Δ_DM_ and measured Δ_DM_, very close to the 1:1 line, with an *r*^2^ larger than 0.85 in all treatments (Fig. 5). Given that *e*_resp_ was determined at plant-scale, we also used a commonly used leaf *e*_resp_ value of –4‰ to calculate *e*_exp_ but has no influence on our conclusions (see discussion below). The fact that there is a very small scattering under each condition strongly suggests that there was also little variation in *e*_exp_ across leaves and plants under a given N/VPD condition.

### Differences between δ^13^C of organs

Δ_ML_ was positively correlated to Δ of different plant organs, and Δ_ML_ was 0.92‰, 0.15‰ and 0.17‰ higher than Δ_Root_, Δ_stem_ and Δ_DT_, respectively (Fig. 6); that is, mature leaves were ^13^C depleted compared with other organs. Moreover, this ^13^C depletion in mature leaves relative to other organs was more significant at high N compared with low N, in line with the larger *e*_exp_ value at high N (Table 2, Fig. 6).

**Fig. 6.**
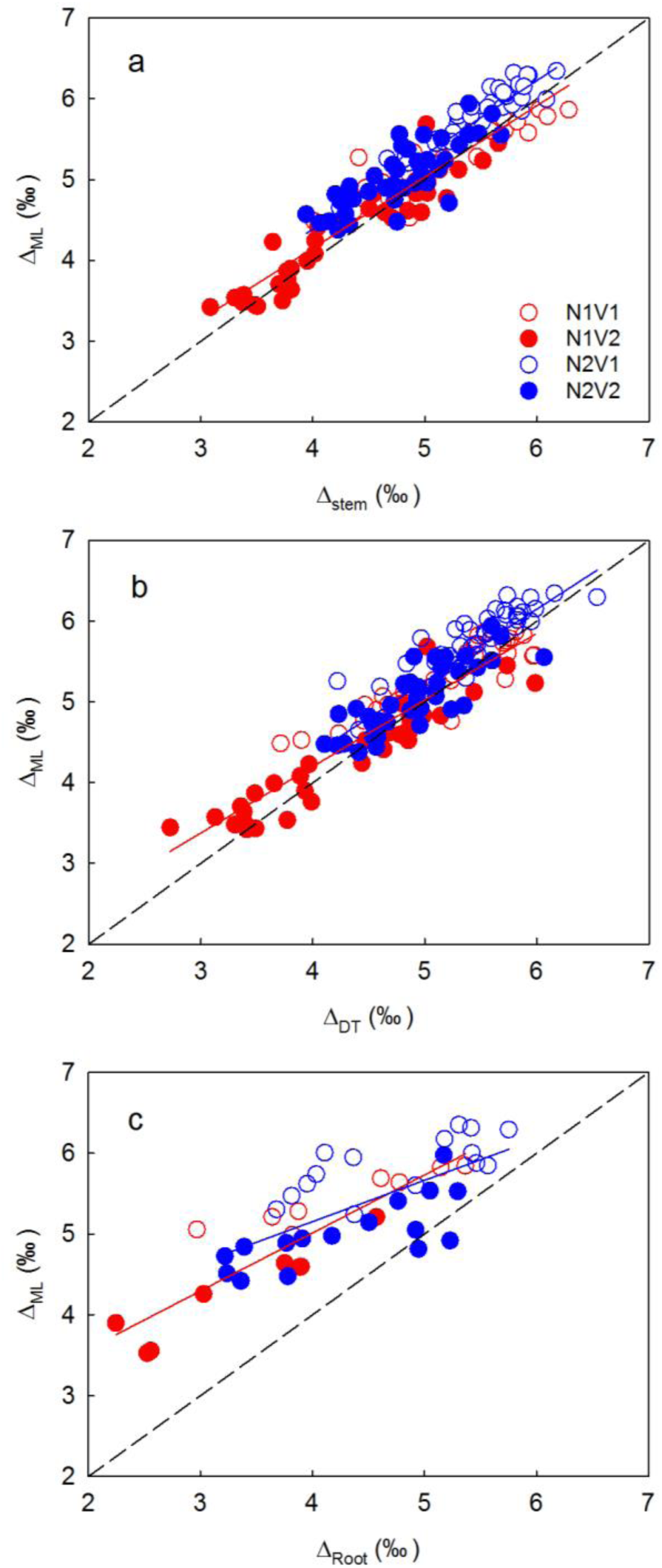
Comparison of Δ_ML_ (mass-weighted Δ of mature leaves on a major tiller) and Δ_stem_ (a), Δ_DT_ (b) and Δ_root_ (c) across N and VPD levels. Red lines are the regression lines of N1 treatment and blue lines of N2 treatments, and the data of low and high VPD of the same N level were pooled for regressions. The dashed line is the 1:1 line for comparison.

## DISCUSSION

### Ontogenetic effect on leaf δ^13^C

Here, we took advantage of the sequential development pattern of leaves to assess the influence of post-photosynthetic fractionations in a C_4_ species. We observed a clear pattern of depletion of leaf organic matter (decreasing δ^13^C value) during ontogeny (Fig. 2a, b), while the δ^13^C of leaf cellulose was unchanged between leaf age categories. This pattern was reproducible when CO_2_ sources with contrasting δ^13^C were used to grow plants. Furthermore, chasing changes in Δ_DM_ of certain young leaves (Nr. 2, 3, and 4) sampled at different dates also show a depletion of leaf material in ^13^C during ontogeny (Fig. S3). Similar results have been observed in field studies across a large geographic region (Fig. S4). Collectively, this indicates that in a C_4_ grass such as *Cleistogenes squarrosa*, leaf δ^13^C declines during ontogeny in both artificial and natural environments. Studies have shown that δ^13^C of young, expanding leaves have less negative values compared with mature leaves in a range of species (Ghashghaie *et al*. 2016; Vogado *et al*. 2020; Gessler *et al*. 2008). While several explanations can be put forward (including a change in photosynthetic properties with leaf age and rank), our results suggest that the decline in leaf δ^13^C during ontogeny is mostly due to post-photosynthetic fractionations, since we found little change in δ^13^C of structural compounds coming from source substrates (here, cellulose).

### Mechanisms behind the depletion of leaf δ^13^C during ontogeny

In principle, three mechanisms could explain the ontogenetic effect on leaf δ^13^C: *(i)* the isotope fractionation during leaf respiration in favour of ^13^C, leaving ^13^C depleted substrates behind; *(ii)* a progressive enrichment in leaves formed sequentially, due to a fractionation in favour of ^13^C during export metabolism; *(iii)* changes in photosynthetic properties (i.e., in isotope fractionation associated with net photosynthesis) of mature leaves once their structural matter has been synthesised.

In fact, hypothesis *(i)* is likely. We found a plant-scale dark respiratory fractionation (*e*_resp_) of about –0.9‰ (Table 2), which is close to the reported *e*_resp_ of C_4_ leaves (range from –0.2‰ to – 1.9‰), significantly lower than that of C_3_ leaves (*c*. –4‰) (Ghashghaie *et al*. 2016; Rooney 1988; Sun *et al*. 2012; Ghashghaie & Badeck 2014). Leaf respiration is the main component (>50%) of the whole-plant respiration of many herbaceous species (Atkin, Scheurwater & Pons 2007; Lehmeier *et al*. 2010). Therefore, the plant-scale *e*_resp_ should be close to the leaf-level respiratory fractionation (Bathellier *et al*. 2008; Klumpp *et al*. 2005). Nonetheless, different *e*_resp_ values of leaf and plant respiration have been reported, resulting from the opposite signal of the respiratory fractionation in shoot (negative) and root (positive) (Schnyder & Lattanzi 2005). That is, leaf-level *e*_resp_ could be lower than –0.9‰. Although leaf-scale *e*_resp_ should be more suitable for our calculations, the *e*_resp_ of single leaf of *C. squarrosa* with small areas cannot be accurately measured using the on-line ^13^C discrimination system that we have (Gong *et al*. 2015), because of small CO_2_ concentration and δ^13^C differences between the air entering and leaving the leaf chamber.

Hypothesis *(ii)* is also likely, since we found that *e*_exp_ was systematically negative (fractionation against ^12^C), with an average value of –0.68‰. That said, if *e*_resp_ was different from the value utilised in calculations, this would have impacted on *e*_exp_. In other words, our estimation of *e*_exp_ could have been influenced by imputing parameters like *e*_resp_ (and the export rate). To address this issue, we looked at the sensitivity of estimated *e*_exp_ with respect to *e*_resp_ and export rate. If an *e*_resp_ of –4‰ was used, we obtained a mean *e*_exp_ of –0.41‰ (Table S1). If we manipulate export rate by ±40%, we obtained a mean *e*_exp_ of –0.28‰ and –1.11‰, respectively (Table S1). That is, our finding of a negative *e*_exp_ (likely within the range of –1.6 to –0.1) of mature C_4_ leaves is robust. So far, the direct assessment of *e*_exp_ is unavailable because measuring δ^13^C of canopy photosynthesis and extracting sucrose are very challenging. Collectively, our results point toward an apparent fractionation in favour of ^13^C during export. This would agree with the finding that sugars transported by phloem sap are enriched in ^13^C compared to leaf soluble sugars (Bögelein, Lehmann & Thomas 2019) and photosynthetically fixed carbon (Gimeno *et al*. 2021). However, Gessler *et al*. (2008) and Lamade *et al*. (2016) showed that differences in leaf-phloem sugars were not significant. Unfortunately, there are no analogical reports for C_4_ species, limiting opportunities for direct comparison. It should be remembered that sucrose transport itself is unlikely to fractionate (the reduced mass effect of isotopic substitution on passive and active transport is negligible); if transported sugars are ^13^C enriched, this is more likely due to a metabolic effect, where sugar interconversions between sucrose, glucose and fructose are associated with isotope effects.

Under our conditions, hypothesis *(iii)* appears unlikely. In principle, the depletion of leaf δ^13^C during ontogeny could be associated to changes in photosynthetic δ^13^C of the leaf after full expansion (i.e., post-maturation isotopic changes). This has been suggested in mature leaves which are ^13^C depleted relative to growing dicot leaves in Vogado *et al*. (2020). To test this mechanism, we compared the photosynthetic discrimination (Δ_A_) measured by combined gas-exchange and isotopic measurements of young and old mature leaves in the same experiment, and published previously (Gong, Schäufele & Schnyder 2017). The results showed that Δ_A_ was not significantly different between young and mature leaves (Table S2), and δ_A_ was slightly higher in old leaves (although being statistically non-significant) compared to young leaves. Therefore, the observed trend of increasing Δ_DM_ (^13^C depletion in organic matter) during aging cannot be explained by the change in δ^13^C of the photosynthetic carbon of a mature leaf.

Also, our mass balance model which took into account respiration and export (hypotheses *(i)* and *(ii)*) was sufficient to estimate leaf Δ_DM_ satisfactorily. Using treatment-specific mean parameters of *e*_resp_ and *e*_exp_, the model explained more than 85% of variation in Δ_DM_ of over 300 mature leaves sampled in each treatment (Fig. 5). The unexplained variance (between 8 and 15%) may be due to the lack of leaf-specific respiration and carbon export rates, or to losses of carbon non accounted for (such volatile organics, sugar and amino acid root exudation, etc.). We recognise that in our model (Eqn 13), the relative contributions of the two key mechanisms (respiration, export) could not be distinguished confidently, since the estimate of *e*_exp_ depended on *e*_resp_. Improving the post-photosynthetic fractionation model would rely on a deeper understanding of mechanisms associated with these processes. So far, *e*_resp_ was mainly measured on leaf/plant placed in darkness although Tcherkez *et al*. (2011, 2017) showed that metabolic pathways and δ^13^C of respired CO_2_ are different between light and dark periods in C_3_ plants. Currently, the isotope fractionation associated with day respiration and its effect on leaf δ^13^C during ontogeny is unknown and should be studied in future work.

### Differences in δ^13^C between autotrophic and heterotrophic organs and treatments

Our study showed significant differences in δ^13^C between autotrophic and heterotrophic organs, namely, δ^13^C of mature leaves was more depleted compared with pseudostems (by –0.17‰), daughter tillers (–0.15‰) and roots (–0.92‰) (Table 2). This finding also agrees with previous reports of C_4_ plants (Badeck *et al*. 2005; Ghashghaie *et al*. 2016; Sun *et al*. 2012), suggesting that mechanisms similar to that associated with respiration and export/import (*e*_resp_ and *e*_exp_) were also operating in other organs. Also, it is believed that ^13^C-enriched transported sugars (such as sucrose) are the main carbon source for heterotrophic organs while other ^13^C-depleted compounds remain in the leaves (e.g., lipids and lignin) (Tcherkez, Ghashghaie & Griffiths 2007; Bögelein, Lehmann & Thomas 2019; Bowling, Pataki & Randerson 2008; Gessler *et al*. 2008; Kodama *et al*. 2008). Another probable reason is that CO_2_ respired from roots tends to be depleted in ^13^C relative to that from leaves (Sun *et al*. 2012; Ghashghaie *et al*. 2016; Ghashghaie & Badeck 2014; Schnyder & Lattanzi 2005). That is, *e*_resp_ of roots could be higher than that of leaves, or even be positive (Schnyder & Lattanzi 2005; Bathellier *et al*. 2008). Taken as a whole, our results indicate that differences in δ^13^C between autotrophic and heterotrophic organs are also related to “post-photosynthetic” fractionation (i.e., associated with respiration and sugar transport).

### Is total organic matter a good proxy for photosynthetic parameters?

It is common practice to use leaf δ^13^C to infer photosynthetic parameters in C_4_ plants, such as bundle sheath leakiness. In fact, the isotope fractionation by net photosynthesis, Δ, in C_4_ leaves is mostly dependent on bundle-sheath leakiness (denoted as *Φ*) and *C*_i_/*C*_a_ according to the model of Farquhar *et al*. (1983). Applying this model, high VPD and high N supply would indicate an increase in *Φ*, as revealed by combined measurements of gas exchange and Δ_A_ (Gong, Schäufele & Schnyder 2017). Accordingly, Δ_DM_ of young and old mature leaves (Δ_DM2_ and Δ_DM10_) and *Φ* calculated therefrom increase at high VPD and N supply. However, due to the difference in Δ_DM_ of about 1‰ between leaf Nr. 2 and 10, estimated *Φ* differed by 40% (Table S3). That is, leaf age is confounding factor when estimating *Φ* from Δ_DM_.

The almost constant Δ_cel_ in all leaf categories suggests that cellulose is probably better to estimate *Φ* since it indicates near-constant assimilation- and export rate-weighted sucrose δ^13^C-signal of source leaves on a mature tiller. As expected, it reflects almost constant leaf physiological status under our constant growth conditions. Furthermore, there were similar responses of Δ_DM_, Δ_cel_ and Δ_A_ and the respective *Φ* values to VPD and N treatments, suggesting that Δ_cel_ represents the isotopic signatures of photosynthetic products and effectively indicates leaf physiological responses to environmental changes. In particular, Δ_cel_ is not affected by leaf age. Δ_cel_ of successive leaves thus could be used as a ’space for time’ records of the efficiency of the photosynthetic apparatus, with the knowledge of the leaf appearance interval and duration of leaf expansion (Yang *et al*. 2016; Liu *et al*. 2017b).

Having said that, there were isotopic offsets between Δ_cel_, Δ_A_ and Δ_DM_ when compared at the same leaf age (Figs. 2-4). In principle, Δ_cel_ should be impacted by enzymatic and positional isotope effects, as the substrate for cellulose synthesis is produced by conversion of sucrose to UDP-glucose via enzymes such as invertase, which fractionates against ^13^C during sucrose cleavage, by 1‰ (Verbančič *et al*. 2018; Gilbert *et al*. 2012; Mauve *et al*. 2009). Hence, although Δ_cel_ appears to be a more robust isotopic proxy to estimate physiological parameters such as *Φ*, a deeper knowledge of relationships between Δ_cel_ and Δ of primary photosynthetic products (i.e., newly synthesized sucrose) is required, including kinetic isotope effects of sugar-converting enzymes.

### Conclusions and perspectives

Here, we showed an age-dependent divergence in isotope composition, between cellulose and total biomass in successive leaves of a C_4_ grass species (*C. squarrosa*). The progressive depletion in δ^13^C of mature leaves is mainly explained by post-photosynthetic fractionation, rather than variations in photosynthetic δ^13^C of source leaves. The influence of post-photosynthetic fractionation was cumulative and thus increased over time, leading to age-dependent patterns in Δ_DM_. Therefore, using a constant, empirical value of post-photosynthetic fractionation Δ_post_ to correct for the difference between photosynthetically-fixed C and leaf matter is probably not adapted to C_4_ grasses with a sequential growth pattern. Although the post-photosynthetic fractionation was less than 2‰ in this study, it is nonetheless significant (in particular in C_4_ species where the photosynthetic fractionation is low) and should be taken into account when applying the model of Farquhar & Cernusak (2012) to calculate other parameters like bundle-sheath leakiness. For this reason, cautions must be taken when using Δ_DM_ to estimate physiological parameters when applying the Farquhar model. Attention should also be given when using data of leaf Δ_DM_ obtained from mixed samples of different leaf ages. According to the present work, Δ_DM_ of the young leaves such as Δ_DM2_ is a better indicator of variation in bundle-sheath leakiness or intrinsic water use efficiency because Δ_DM2_ is less impacted by the cumulative effect of post-photosynthetic fractionation. Alternatively, Δ_cel_ of leaves is a more reliable physiological proxy as it is similarly sensitive to changes in N nutrition and VPD but not appear to be affected by respiration and export associated fractionation after maturity of the leaves. Still, we recognise that future work should focus on the δ^13^C difference between photosynthetically-fixed carbon and leaf cellulose to enable quantitative prediction of photosynthetic parameters from leaf cellulose δ^13^C. Also, more compound-specific δ^13^C analyses (via GC-c-IRMS or LC-co-IRMS) of C_4_ leaves and phloem sap would be useful to determine metabolic mechanisms of post-photosynthetic isotope fractionations.

## ACKNOWLEDGMENTS

This work was supported by the National Natural Science Foundation of China (NSFC 32120103005, 31870377 and 31901090) and the Deutsche Forschungsgemeinschaft (SCHN 557/7-1 and SCHN 557/9-1).

## AUTHOR CONTRIBUTIONS

HS, XYG and RS designed and planned the research; HTL, FY, LL and RS performed the sampling and isotope analyses; YZY analysed the data; XYG and GT established the isotopic mass-balance equations; YZY and XYG wrote the manuscript and all authors contributed to the revision.

---

**Table S1.** Sensitivity test on *e*_exp_ by varying the *e*_resp_ (observed values and *e*_resp_=-4‰) and export rate (60% and 140%). Values are means ± SE (*n*=8).

| Treatments | $e_{\text{exp}}$ | | | |
| --- | --- | --- | --- | --- |
| | $e_{\text{resp}}=\text{observed}$ | $e_{\text{resp}}=-4\text{‰}$ | 60% export rate | 140% export rate |
| N1 VPD1 | $-0.57 \pm 0.06$ | $-0.23 \pm 0.04$ | $-0.86 \pm 0.12$ | $-0.19 \pm 0.05$ |
| N1 VPD2 | $-0.50 \pm 0.06$ | $-0.30 \pm 0.04$ | $-0.83 \pm 0.09$ | $-0.36 \pm 0.04$ |
| N2 VPD1 | $-0.96 \pm 0.04$ | $-0.67 \pm 0.03$ | $-1.60 \pm 0.07$ | $-0.38 \pm 0.08$ |
| N2 VPD2 | $-0.68 \pm 0.04$ | $-0.45 \pm 0.03$ | $-1.14 \pm 0.06$ | $-0.18 \pm 0.04$ |

**Table S2.** Carbon isotope discrimination during net CO_2_ exchange (Δ_A_) of young and old leaves of *C. squarrosa* grown under low or high N supply combined with low or high VPD. Data were shown as mean ±SD (*n*=3-4). The irradiance during measurements was 1500 μmol m^-2^ s^-1^; average leaf temperature was 27°C; and average CO_2_ concentration in the leaf chamber was 380 μmol mol^-1^. Measurement details were shown in Gong *et al*. 2017.

|  | N1 |  | N2 |  |
| --- | --- | --- | --- | --- |
|  | VPD1 | VPD2 | VPD1 | VPD2 |
| Leaf Nr. 3 | 5.09 ± 0.64 | 5.67 ± 1.31 | 5.20 ± 2.32 | 5.94 ± 0.68 |
| Leaf Nr. 7 | 4.02 ± 2.10 | 4.60 ± 1.65 | 5.36 ± 1.34 | 5.84 ± 1.32 |

**Table S3.**
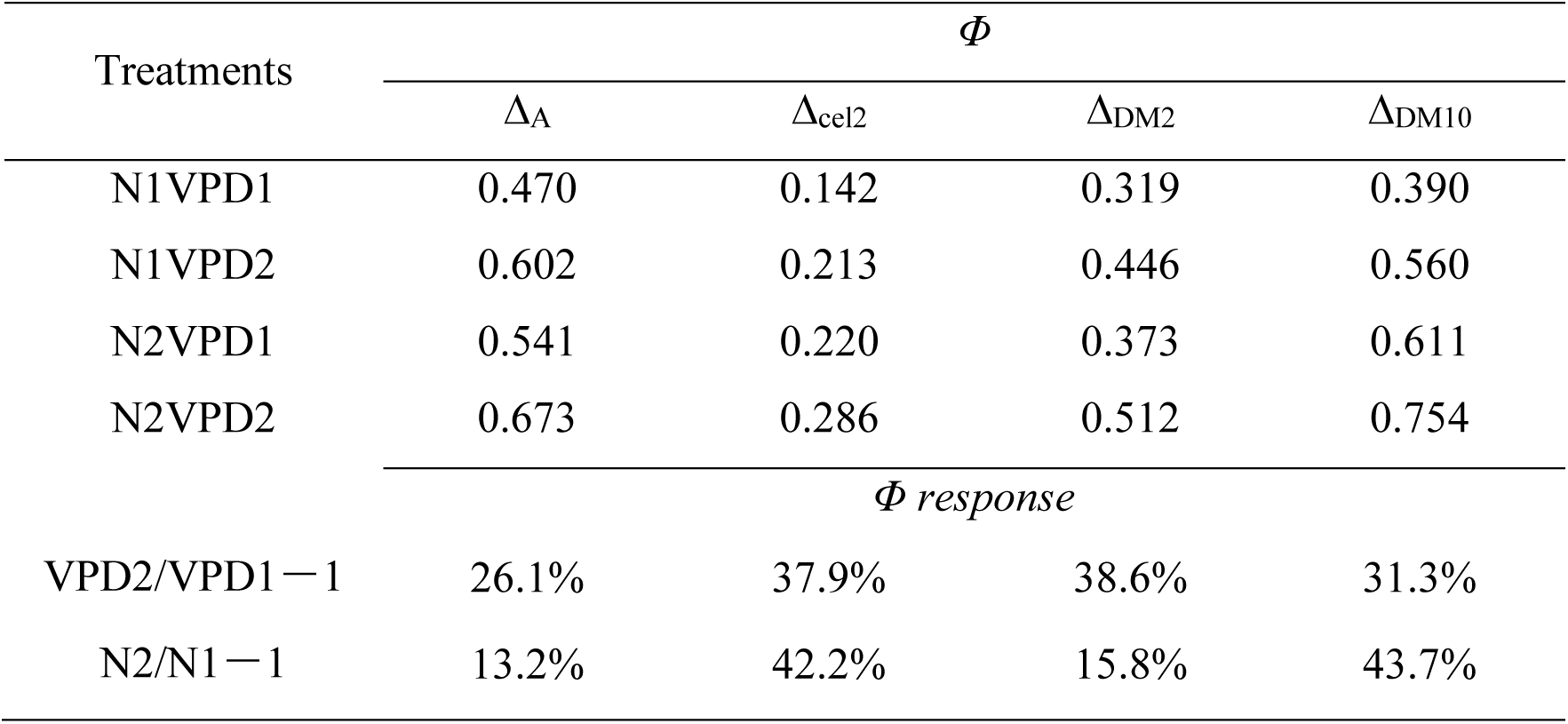
Bundle sheath leakiness (*Φ*) of *C. squarrosa* grown under low or high N supply combined with low or high VPD were calculated from on-line measurements (Δ_A_, the results have been published in Gong *et al*. 2017), cellulose of leaf Nr. 2 (Δ_cel2_), dry matter of leaf Nr. 2 (Δ_DM2_) and leaf Nr. 10 (Δ_DM10_). *Φ* in *C. squarrosa* was estimated by the simple model of C_4_ carbon isotope discrimination (Farquhar 1983) as Δ = *a* + (*b*_4_ + *Φ* (*b*_3_ – *s*) – *a*) *C*_i_/*C*_a_, where *a* is the fractionation during diffusion of CO_2_ in air through stomata (4.4‰), *b*_4_ is the fractionation associated with PEP carboxylation and preceding isotopic equilibrium during dissolution and conversion to bicarbonate (–5.7‰ at 25℃), *b*_3_ describes the fractionation by Rubisco (29‰), and *s* is the fractionation during leakage of CO_2_ out of the bundle sheath cell (1.8‰).

**Fig. S1.**
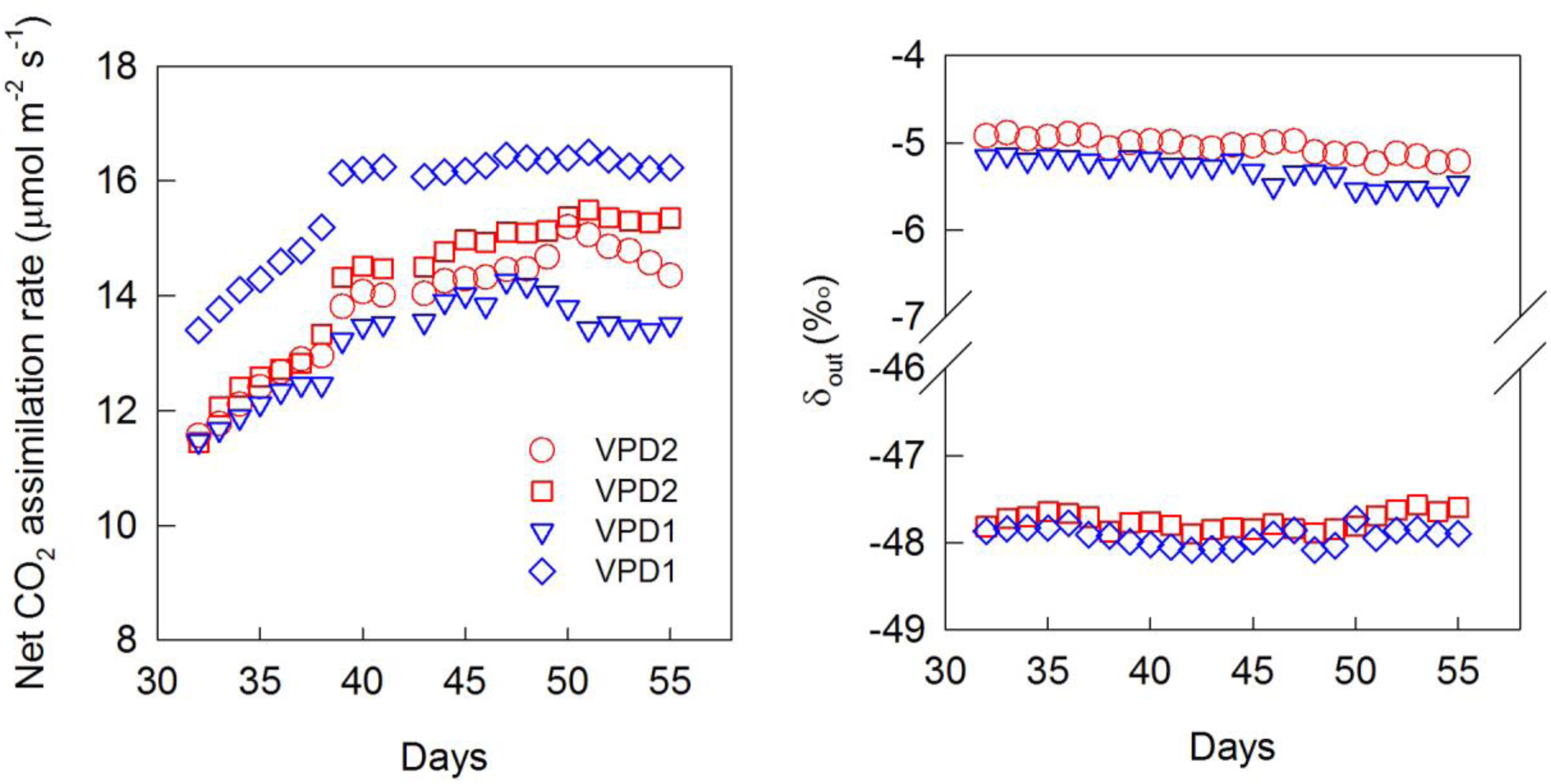
Examples of mesocosm-scale net assimilation rate and δ of the chamber air (δ_out_) of the moderate N experiment (N1) during the experimental period. The symbols represent data of different growth chambers.

**Fig. S2.**
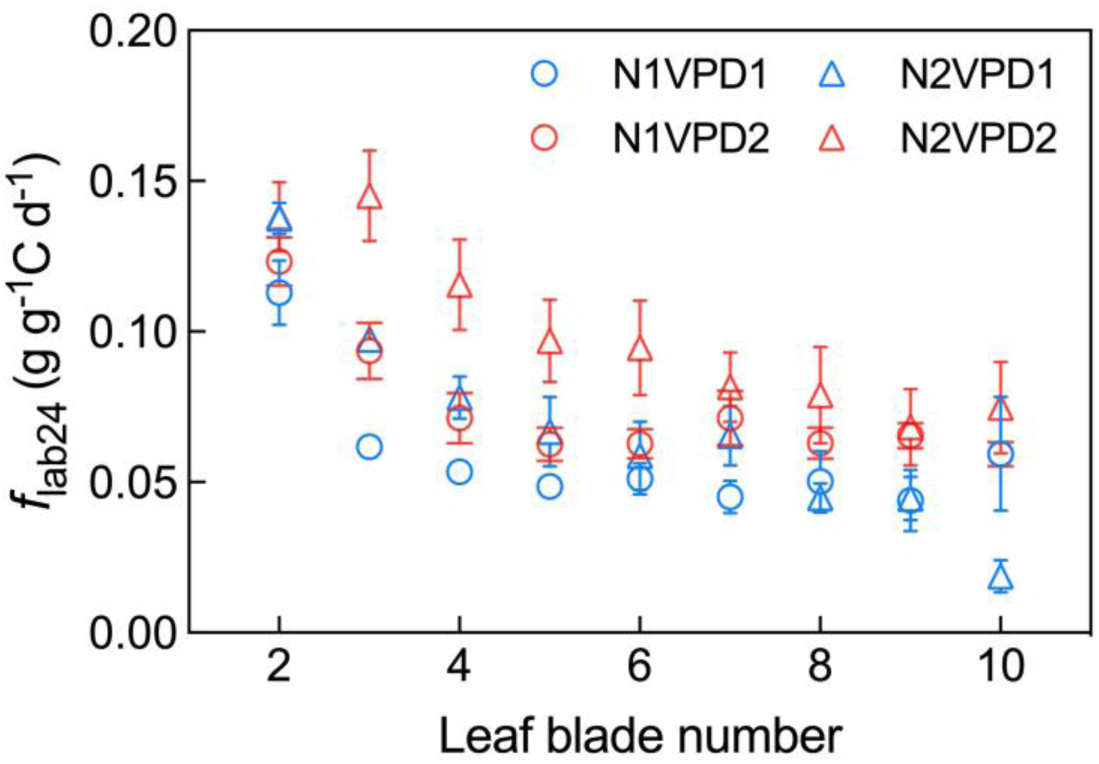
*f*_lab24_ of mature leaves under different N supply and VPD treatments. Figure points are means ±standard error, *n*=2-4.

**Fig. S3.**
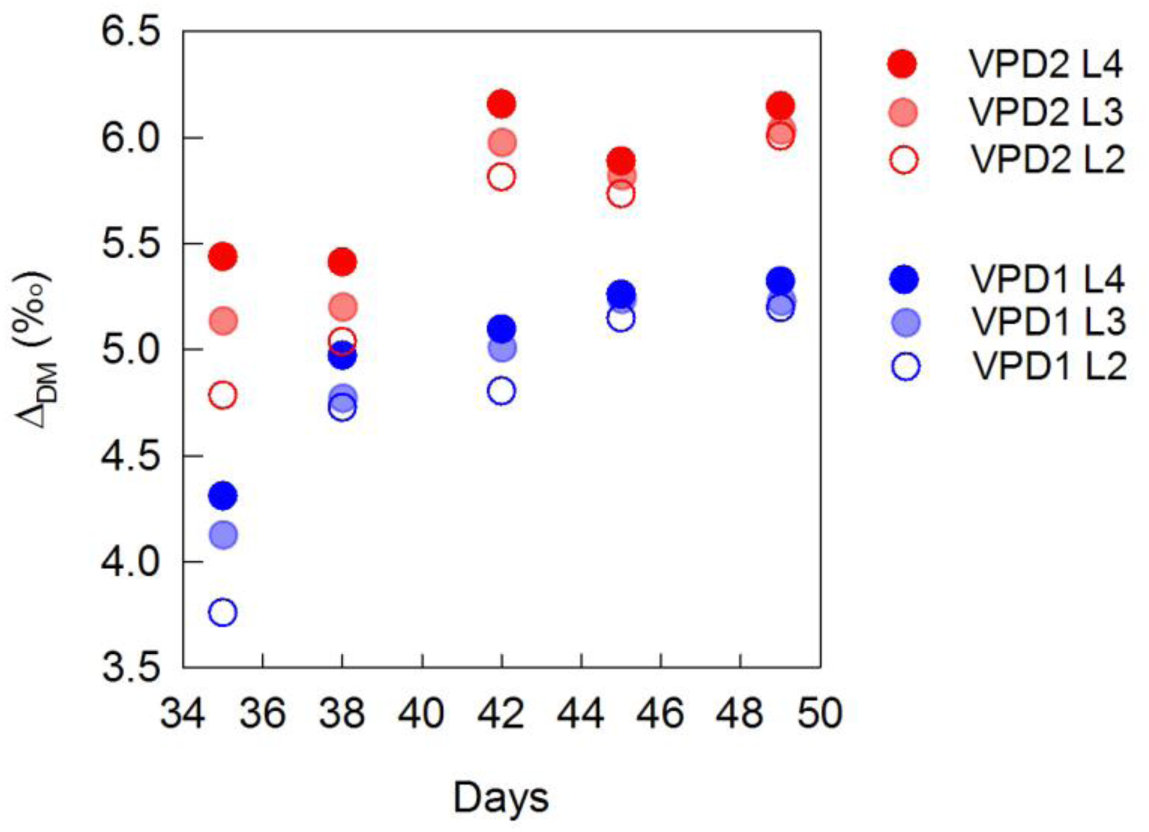
Changes in Δ_DM_ of leaf Nr. 2, 3, and 4 in different sampling dates during ontogeny of the experimental run with moderate N (N1).

**Fig. S4.**
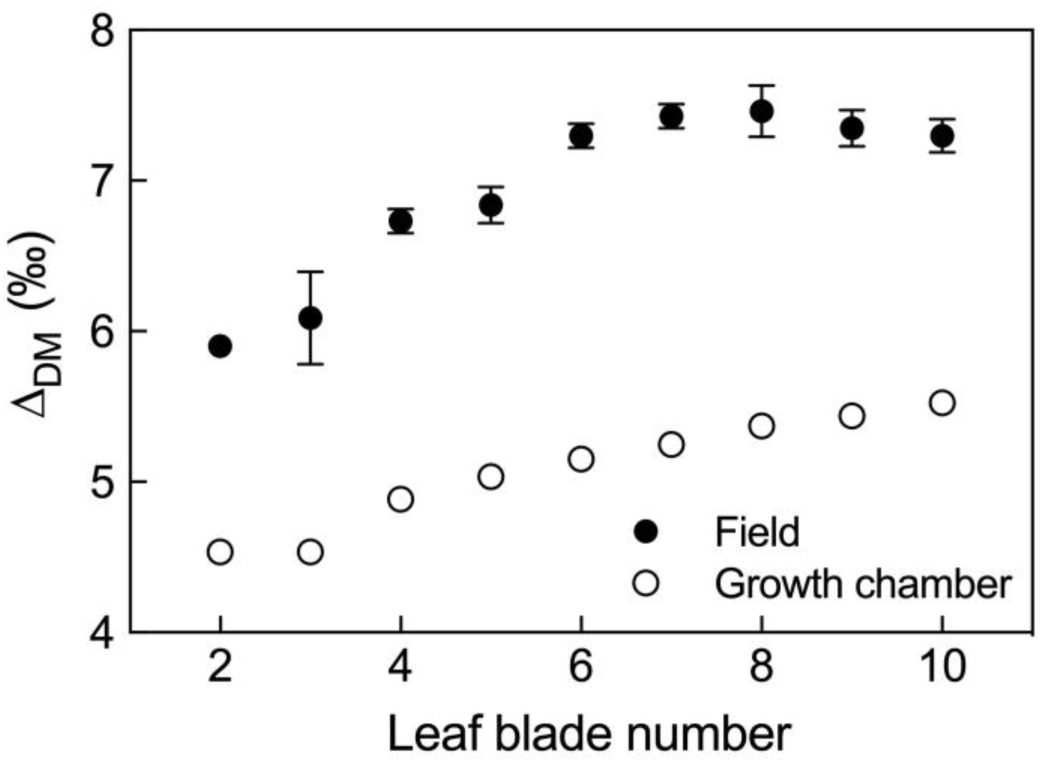
Comparison of observed Δ_DM_ of mature leaves in *Cleistogenes squarrosa* growing in growth chamber and field. Δ_DM_ of field-growth leaves (filled symbols) were collected from Yang *et al*. (2011) and Δ_DM_ in growth chamber (open symbols) were derived in the present study for all treatments. Error bars denote the standard error (SE).

